# Spatial causal discovery reveals novel mechanisms of cellular senescence in human liver

**DOI:** 10.64898/2026.11.13.766422

**Authors:** Erich Kummerfeld, Yinzhao Wang, Leland Williams, Samuel T. Peters, Elizabeth L. Schmidt, Mickayla DuFresne-To, Paul Robbins, Mac Anderson, Sayeed Ikramuddin, Oyedele Adeyi, David Haynes, Steven Johnson, Jinhua Wang, Nuri Alpay Temiz, Laura Niedernhofer, Andrew Nelson, Constantin Aliferis

## Abstract

The complex network of mechanisms that surround cellular senescence, such as its precise causes and effects, are not entirely understood at present. Senescence occurs when a cell permanently ceases dividing, halts apoptosis, stops performing its normal functions, and begins secreting chemicals that can induce other cells in its environment to also become senescent.

Senescent cells are present in both healthy and unhealthy tissue, but their abundance is positively correlated with disease and age. A better understanding of senescence could lead to healthier aging, improved recovery from chemotherapy, or increased lifespan. This study used novel machine learning algorithms - with guaranteed properties of correctness under broad assumptions - to investigate senescence in human liver tissue by analyzing Visium spatial transcriptomics data. This produced data-driven models of the causes and effects of senescence, both locally and from nearby tissue. This report includes an assessment of whether the data meet key requirements for causal analysis, the transcripts identified by causal analyses as causes and effects of senescence marker genes, and pathway analyses of those gene sets.

**Key Takeaways:** 1. Senescence in Visium data from human liver tissue is spatially autocorrelated, exhibiting significant spatial clustering.

2. We show that Visium data from human liver tissue satisfies the causal Markov condition.

3. We introduce novel spatial extensions of established causal discovery machine learning algorithms and apply them on Visium data from human liver tissue. The causal modeling produced lists of transcripts directly (i) upstream and (ii) downstream of senescence marker genes, which had relatively little overlap with standard senescence marker lists SenMayo and SenUMN.

4. Pathway analyses of the upstream and downstream transcripts reveal that senescence-associated signaling integrates cellular stress inputs and immune responses to fuel a self-sustaining loop of integrin-mediated TGF-β activation and continuous matrix deposition.

## Introduction

Senescence is a cellular state where the cell permanently halts the normal cell cycle by preventing the two normal lifecycle outcomes: cell division and apoptosis (programmed cell death). Senescence has been widely observed in humans and many other species, including most mammals and birds, and many invertebrates. Both within and across species, senescence is closely associated with age: younger organisms tend to have fewer senescent cells, and species capable of extreme lifespans also tend to have fewer senescent cells. Further, both within organisms and across species, the abundance of senescent cells is also associated with disease. Overall, it is plausible that a deep understanding of senescence could lead to precise interventions that result in healthier aging or increased lifespan in humans.

At present, much remains unknown about the specific mechanisms of senescence. Senescence can be induced in cells in laboratory conditions, typically by having them repeatedly divide or by exposing them to stress such as high levels of toxicity or radiation[1–4]. Laboratories have also demonstrated that a living organism’s apparent biological age can be increased or decreased by adding or removing senescent cells[5–7]. At the transcriptomic level, many genes have been associated with senescence, in a variety of different animal models and tissue types. Gene lists have been curated to help use transcriptomic data for the identification of senescence in settings where the defining phenotypic markers (cell cycle arrest) can not be observed[8].

Existing knowledge about transcripts and senescence is of two types: (1) a gene may be a known part of a pathway that relates to the senescence phenotype, such as pathways that halt apoptosis or cell division; (2) a gene transcript may have been observed to be present more or less often in a group of senescent cells than in a control group of non-senescent cells, i.e. it may be statistically associated with senescence. Both of these types of knowledge have limitations.

In the first case, there is a clear possible mechanism by which the transcript could possibly be related to senescence. One limitation of this is that there is no guarantee this mechanism is active, or that there aren’t existing biological control mechanisms to counteract this transcript’s potential influence on senescence. Another limitation is that we do not have a complete picture of all biological pathways, and using only known pathways may exclude genes that influence senescence, or are influenced by it, via as-yet unknown pathways.

Elucidating the second case, there is a clear empirical connection between increased or decreased expression of that gene transcript in actual senescent cells. However knowledge of the role that the gene plays, if any, is limited. A transcript could be associated with senescence for any of the following reasons: (a) the gene may play a critical role in producing and sustaining senescence; (b) the gene may promote or suppress other genes that play critical roles in senescence; (c) the gene may be promoted or suppressed by other genes that are critical to senescence; (d) the gene may be promoted or suppressed by other factors that also promote or suppress genes that are critical for senescence. Or in short, knowing that a gene is associated with senescence in cells only tells us that it is likely either part of senescence, a cause of senescence, an effect of senescence, or has a confounded relationship with senescence, or any combination of those.

New sources of information about senescence have recently become available through innovations in measurement and data analysis technologies. Spatial transcriptome measurement technologies such as Visium allow for the investigation of senescence within the context of the surrounding tissue. Causal discovery machine learning methods enable the production of data-driven causal models from data. Applying causal discovery to spatial omics data has the potential to offer insights that are molecular in specificity, mechanistically interpretable, and incorporate spatial context information that is critical for understanding how senescence develops and spreads.

State of the art causal discovery algorithms operate under well understood distributional assumptions that when satisfied ensure correct results. However they are not designed for spatial data. Here we (a) extend leading causal discovery algorithms for the spatial domain and (b) test their assumptions in real biological data. We then (c) use the algorithms to derive novel findings and insights from the data.

Specifically, this study was guided by the following questions:

1. Are senescence biomarkers spatially autocorrelated?
2. Do senescence biomarkers satisfy a spatial causal Markov property?
3. What molecular targets are upstream, and what molecular outcomes are downstream, of senescence signature and biomarker genes?
4. Can the novel gene-level spatial causal discovery approach introduced here produce valuable new insights into cellular senescence?

## Background and Related Work

The National Cancer Institute defines cellular senescence as: “a process by which a cell ages and permanently stops dividing but does not die. Over time, large numbers of old (or senescent) cells can build up in tissues throughout the body. These cells remain active and can release harmful substances that may cause inflammation and damage to nearby healthy cells. Senescence may play a role in the development of cancer and other diseases.” [9] Cellular senescence may hold the keys to longer lives, healthier aging, and disease treatment and prevention [4,10]. While much has been learned, many aspects of senescence remain unknown [11–15].

Like cancer cells, senescent cells derive from previously healthy cells performing normal operations as part of some tissue or organ. Senescent cells from different tissues can have a different character, as well as different implications for age-related diseases. In this paper we investigate data from human liver samples. Previous investigations have also studied senescence in liver, both human and mouse, and associated the prevalence of senescent cells with a variety of liver-specific stress exposures and diseases [16–21].

Spatially, senescent cells are recognized to have distinct impacts on their microenvironment and the surrounding cellular organization. Senescent cells have a distinct secretary profile composed of senescence associated secretary phenotype (SASP) factors. Some prior studies have begun to describe the spatial genomic signatures of senescent cells and their microenvironments, however much remains unknown [11,20,22–25]. Analyses reported in this paper expand on the existing knowledge about the spatial character of senescence by evaluating its spatial autocorrelation and whether it satisfies the Spatial Markov Condition.

Substantial prior work has investigated the possible molecular pathways that induce or are induced by senescence. We highlight a few key papers here. In [26] they identify multiple channels that could lead to senescence, and identify p21 and p16 as the critical pair of genes that together mediate all effects of stress on downstream senescence. The heterogeneity of senescence is identified in many papers, including [27] where the lack of consistent mechanisms or markers is identified as major challenges. p53 and pRB tumor suppressor pathways have also been named as important controllers of cellular senescence [28,29].

Applications of causal discovery and similar methods for learning causal structural models from omics data have been extensive in the non-spatial case, but for spatial omics data they are limited. There has been some progress developing new methods tailored to spatial omics platforms [30,31], however (a) these have not been accompanied by rigorous testing of their modeling assumptions in real data, and (c) they have not been applied in the context of senescence.

Here, we (1) extend existing high performance causal discovery algorithms, that have guarantees of correctness under broad assumptions, to the spatial omics context by constructing spatial lag variables; (2) evaluate whether our available Visium human liver data satisfies the distributional assumptions required to guarantee correctness of the methods; (3) use the novel algorithms to characterize the molecular causes and effects of a panel of previously known senescence marker genes specific to human liver tissue; (4) perform pathway membership and enrichment analyses on these upstream and downstream genes, in order to better characterize the mechanisms of senescence in human liver tissue.

## Methods

### Data

Liver tissue samples from 10 human participants (8 female, 2 male) were analyzed. Ages ranged from 32 to 70. Liver health varied from normal to highly fibrotic, according to pathology evaluation. See Table 1 for a detailed description of the population. Samples 110_A1 and 110_B1 were from the same participant.

**Table 1:** Liver tissue samples, pathology evaluation, and demographics

| **Visium Slide ID** | **SenNet ID** | **Age, Years** | **Sex** | **Macrosteatosis (%)** | **Steatosis Score** | **Ballooning** | **Inflammation** | **Fibrosis** |
|---|---|---|---|---|---|---|---|---|
| V12M15-110_A1 | SNT745.FMZR.774 | 70 | Female | 70 | 3 | 1 | 2 | 1c |
| V12M15-110_B1 | SNT275.PRRJ.325 | 70 | Female | 70 | 3 | 1 | 2 | 1c |
| V12M15-110_C1 | SNT477.ZMNT.864 | 48 | Female | 5 | 0 | 0 | 0 | 0 |
| V42D20-289_A1 | SNT896.QDXS.668 | 42 | Female | 0 | 0 | 0 | 1 | 0 |
| V42D20-289_D1 | SNT696.QGBM.388 | 38 | Female | 1 | 0 | 0 | 0 | 0 |
| V42L06-053_A1 | SNT346.LGHF.822 | 50 | Male | 0 | 0 | 0 | 1 | 4 |
| V42L06-053_D1 | SNT673.JSSW.854 | 37 | Female | 0 | 0 | 0 | 0 | 0 |
| V42L29-055_A1 | SNT782.NQRT.677 | 49 | Male | 0 | 0 | 0 | 0 | 0 |
| V42L29-055_D1 | SNT387.THHP.768 | 50 | Female | 10 | 1 | 0 | 0 | 0 |
| V42L29-278_A1 | SNT467.RFZM.575 | 42 | Female | 20 | 1 | 1 | 1 | 1a |
| V42L29-278_D1 | SNT576.ZVWX.827 | 32 | Female | 0 | 0 | 0 | 0 | 0 |

All tissue samples were imaged using the Visium platform and Cytassist workflow. The workflows for storing, preparing, and imaging these samples has been previously described in [32].

### Data cleaning and standardization

To clean the data, we first applied the artifact removal technique BLADE [32] and removed spots detected as edge artifacts, border artifacts, and location malfunction artifacts for all tissue samples. To control for variation in total reads across the image, the number of gene reads in each spot was normalized to the total reads in that spot. In other words, counts were transformed to gene-specific spot-level proportions. These proportions were then converted to z-scores across all spots in the tissue sample. After this standardization procedure, a value of 0 for some gene X in spot Y from tissue Z indicates that the proportion of gene X among all genes in spot Y is the mean average found across all the spots in tissue Z. More positive values indicate that the concentration of reads for that gene, relative to other spots in the same tissue sample, are greater in that spot; negative values indicate that the concentration is less.

### Senescence signature

We used the Mean of Four Means (M4M) senescence signature as our target gene list. This list has 10 genes in 4 categories: Collagen marker COL1A1, COL1A2 and COL3A1; Senescence markers CDKN1A and JUN; SASP factors GDF15, SERPINE1 and TIMP1; and Anti-apoptosis markers BCL2L1 and MCL1. Since we are interested in specific gene-gene interactions, we did not look at how other genes interact with these aggregate M4M signals but rather inspected how other genes interact with each of the 10 genes used in the M4M signature.

### Gene selection for analysis

To reduce computational complexity, we restricted our analyses to genes that were commonly associated with the M4M genes across our 11 tissue samples. First, feature selection was performed using HITON-PC (with k=3) on each tissue sample separately [33]. This algorithm returns the Markov Boundary of the M4M group, i.e., the genes that could not be made independent of one or more M4M genes even after conditioning on up to 3 genes not in the Markov Boundary. To arrive at a universal list of genes to use for all tissue samples, the intersection of the tissue sample-specific gene lists was used. To confirm the quality of this gene list, root mean squared error of approximation (RMSE) was calculated for predicting the M4M genes from the final feature list.

### Spatial autocorrelation or clusteredness

To assess whether spatial context needed to be accounted for, rather than treating Visium spots as independent observations, global Moran’s I was calculated for every gene in the selected gene set as well as the 10 target genes. For each gene, expression values across all spots were tested for positive spatial autocorrelation using “spdep::moran.test()” function from R. Under the randomization null hypothesis of no spatial association between gene expression and spatial location, the expected value and variance of Moran’s I were calculated from the spatial weights matrix derived from Visium spot spatial arrangement. The observed Moran’s I was then standardized, and the p-value was obtained from the standard normal distribution using a one-sided test for positive spatial autocorrelation. Thus, the test is a normal-approximation (z) test of Moran’s I under the randomization null, rather than an explicit permutation test.

### Constructing spatial lag variables

Our primary spatial analyses calculate spot-by-spot statistics, with spots as the unit of analysis. In order to capture the *spatial context* information of each spot, spatial lag variables were constructed. These consist of additional variables that summarize other spots that are located in specified ways to the primary spot that defines each record. Let Y be an arbitrary spot on a Visium slide. Let Yd be the average value of the transcripts in all available spots that are at d hexagonal taxicab distance from Y. So for example, Y0 is the value within the spot itself, while Y1 is the average value of all neighboring spots, and Y2 is the average value of spots just surrounding the neighbors, and so on. This forms a table with n rows, where n is the number of spots on the slide, and m+1 variables, where m is the maximum distance that is considered. In this table, variable Y0 indicates the values of the spot that defines the row, and variables Y1 through Ym are the spatial lags which capture the surrounding values.

### Validity of spatial causal discovery: testing the Spatial Causal Markov Property

The Causal Markov Property (CMP) is a fundamental assumption for causal discovery algorithms with guaranteed correctness [34]. It is typically assumed on grounds of the nature of macroscopic causal systems, but is not easy to test in practice. In the context of our work we define a spatially-specific version of the CMP which we call the Spatial Causal Markov Property (SCMP).

SCMP is satisfied for distance D if on a collection of spatially located variables if: the value of any variable X with location L is statistically independent of any variable Z at a distance greater than D from L, conditional on the values of variables at all locations with distance less than D from L. This is a special case of the more general Spatial Markov Property [35,36]. The intuition behind this property is that any causal relationship between spatially separated transcripts is expected to be completely mediated by the physical materials that lie between them. Importantly, unlike the more general CMP, the SCMP is testable in real world data.

To test the SCMP on our Visium data, for each distance d = 2,…,6, we tested the conditional association between Y0 and Yd while controlling for all intermediate layers Y1,…,Yd−1. Specifically, we (a) fit the linear model Y0 = β0 + β1Y1 + … + βd−1Yd−1 + βdYd + ε; and then (b) tested the null hypothesis H0: βd = 0. If βd is not significantly different than 0, then this implies that Y0 ⟂ Yd | Y1,…,Yd−1. As such, if H0 is accepted for some value of d, then the data satisfy the SCMP at that distance, since by construction the variables Y1,…,Yd-1 spatially separate Y0 from Yd. We record the smallest value of d that satisfies SCMP for each M4M gene on each tissue sample.

### Causal discovery modeling

There are hundreds of causal discovery algorithms that have been developed (and infinitely large families have been characterized), and a large body of work has validated them in controlled conditions and real life research (e.g. [37]). Prior algorithms were not designed for spatial data so here we pursued spatial extensions. We selected two algorithm families that (a) are published and have well established properties including guarantees of correctness and completeness, (b) have shown exceptional performance in past studies, and (c) are operationally dissimilar, with distinct theoretical and practical features. Specifically, they are the Best Order Score Search (BOSS) [38] algorithm and the Local-to-Global-Learning (LGL) [39] algorithm family. Both algorithms can take in tabular observational data, such as spatially-lagged Visium data, and return graphical models with causal interpretations. BOSS uses a score-based approach that searches through the space of causal orderings of the variables to find a DAG that maximizes goodness of fit. LGL uses a multiple-testing approach to first learn local Markov blankets for each variable [40] using the LGL family of local causal algorithm discovery and then integrates those local neighborhoods to form a global network.

We call the spatial version of LGL, LGLsrsi (standing for LGL, spatially resolved, spatially invariant) and the spatially resolved BOSS, BOSSsrsi. LGLsrsi and BOSSsrsi, extend LGL and BOSS by introducing: (i) an assignment of explicit spatial information to each variable; (ii) an extension of theory of the base algorithms (LGL, BOSS) to produce sound and complete spatial causal models under sufficient assumptions, (specifically the SCMP, explained before, and the spatial invariance assumption (SIA); the SIA requires that the local causal interactions hold the same regardless of where in space they occur (which in spatial analytics corresponds to the second-order spatial invariance models); (iii) practical tests for testing the assumptions of correctness/completeness; (iv) data pre-processing and normalization steps specific to spatial analysis and the Visium platform specifically; (v) a protocol for synthesizing and selecting information from models built from different biological samples. We used the recommended default tuning parameter values for both algorithms. All searches were performed using computing resources at the Minnesota Supercomputing Institute.

### Identifying genes upstream and downstream from senescence

The causal discovery analyses yield structural models for each tissue sample. Variation among tissue samples, whose models can contain hundreds or thousands of gene-gene relationships, thus becomes necessary. The following steps were used to synthesize and select information from this collection of models, and are performed separately for BOSSsrsi and LGLsrsi:

1. Calculate which genes were most commonly modeled as directly *upstream* from the M4M genes. For every edge in a model pointing from a non-M4M gene to an M4M gene, add the non-M4M gene to a list. Count the number of times this happens (node counts) for each of the non-M4M genes. For the following GSEA and Pathway analysis, use all genes in the list that have a total count of 3 or more across all models.
2. Calculate which genes were most commonly modeled as directly *downstream* from the M4M genes. For every edge in a model pointing from a M4M gene to a non-M4M gene, add the non-M4M gene to a list. Count the number of times this happens (node counts) for each of the non-M4M genes. For the following GSEA and Pathway analysis, use all genes in the list that have a total count of 3 or more across all models.
3. Compute the number and percentages of genes in each list that do and do not occur in the SenMayo or SenUMN lists of senescence signature genes.
4. Perform gene set enrichment analysis (GSEA) and Pathway analysis, described next, on each of these sets of genes.

### Pathway Analysis

Genes found to be upstream of M4M or downstream of M4M were used with node counts as “expression” values. GSEA and pathway over representation analysis was performed using the R shiny app TranscriptoPathR (github.com/temizna/TranscriptoPathR) and ClusterProfiler [41].

## Results

### Here we report findings from applying the above methods to our samples of human liver Visium data

#### Data cleaning and pre-processing

After cleaning the data with BLADE, the 11 tissue samples had a total of 35,385 spots, averaging approximately 3,217 spots per tissue sample with the minimum being 1,172 spots and the maximum being 4,572 spots. The specific number of spots available per tissue sample are available in Appendix A.

### After running the gene selection process, 2,558 genes were selected for analysis

#### Spatial autocorrelation

Among all the selected non-M4M genes, 97.8% rejected a Moran’s I null hypothesis test of no spatial autocorrelation (p < 0.05), indicating significant positive spatial autocorrelation. Among all M4M genes, 100% rejected Moran’s I null hypothesis tests (p < 0.05).

#### Spatial Causal Markov Property

All 10 M4M genes were tested on all 11 tissue samples. This resulted in 110 tests, out of which 10 satisfied the Spatial Markov Property at a distance of 3 and the remaining 100 satisfied the Spatial Markov Property at a distance of 2. The complete table showing the minimum distances that satisfy the Spatial Markov Property for all gene-sample pairs, and the corresponding p-values, is in Appendix B.

#### Causal discovery

Table 2 shows the total counts and overlap of the genes modeled as directly upstream and downstream of the target senescence genes by BOSSsrsi and LGLsrsi. In total, BOSSsrsi yielded 224 directly upstream genes and 659 directly downstream genes. LGLsrsi yielded 38 directly upstream genes and 53 directly downstream genes, reflecting different inductive bias tradeoffs or parsimony vs completeness in the small sample. Tables 3, 4, 5 and 6 show the top genes by count from each list, along with whether that gene is part of the SenUMN or SenMayo lists. The complete gene lists, and corresponding tables with counts and additional information, are available online (https://github.com/KummerfeldLab/Visium_causal_results). The genes discovered through this process have varying degrees of overlap with the SenUMN and SenMayo senescence signature gene lists. BOSSsrsi identified much larger gene sets than LGLsrsi, and few of them appeared in the SenUMN or SenMayo gene lists: the largest overlap is SenMayo identifying 13.4% of the genes in the Upstream BOSSsrsi list. The LGLsrsi method produced relatively fewer genes than BOSSsrsi, with 47.4% of its upstream genes and 32.1% of its downstream genes appearing in the SenMayo list, respectively. SenUMN, being a smaller list than SenMayo, also had smaller overlap with all lists.

**Table 2:** Upstream and Downstream Genes Count and Overlap

| Method | Upstream or Downstream | Total count | SenUMN overlap # | SenUMN overlap % | SenMayo overlap # | SenMayo overlap % |
|---|---|---|---|---|---|---|
| BOSSsrsi | Upstream | 224 | 7 | 3.1% | 30 | 13.4% |
| BOSSsrsi | Downstream | 659 | 2 | 0.3% | 23 | 3.5% |
| LGLsrsi | Upstream | 38 | 4 | 10.5% | 18 | 47.4% |
| LGLsrsi | Downstream | 53 | 4 | 7.5% | 17 | 32.1% |

**Table 3:** BOSSsrsi top 12 directly upstream genes

| In SenUMN? | In SenMayo? | **Gene** | **Total** |
|---|---|---|---|
| FALSE | TRUE | IGFBP7 | 43 |
| FALSE | FALSE | DCN | 37 |
| FALSE | FALSE | TMSB4X | 36 |
| FALSE | FALSE | MGP | 34 |
| FALSE | FALSE | FOS | 32 |
| TRUE | TRUE | IGFBP3 | 32 |
| FALSE | FALSE | CD74 | 28 |
| FALSE | FALSE | PTGDS | 23 |
| FALSE | FALSE | JUNB | 21 |
| FALSE | FALSE | VIM | 21 |
| FALSE | FALSE | LYZ | 21 |
| FALSE | FALSE | B2M | 21 |

**Table 4:** BOSSsrsi top 12 directly downstream genes

| In SenUMN? | In SenMayo? | **Gene** | **Total** |
|---|---|---|---|
| FALSE | FALSE | COL6A3 | 39 |
| FALSE | FALSE | COL6A2 | 34 |
| FALSE | FALSE | COL6A1 | 30 |
| FALSE | FALSE | TNXB | 26 |
| FALSE | FALSE | SPARC | 25 |
| FALSE | TRUE | MMP2 | 24 |
| FALSE | FALSE | THY1 | 21 |
| FALSE | FALSE | DCN | 20 |
| FALSE | FALSE | DPT | 20 |
| FALSE | FALSE | COL4A1 | 19 |
| FALSE | FALSE | MXRA8 | 19 |
| FALSE | FALSE | FSTL1 | 18 |

**Table 5:** LGLsrsi top 11 directly upstream genes

| In SenUMN? | In SenMayo? | **Gene** | **Total** |
|---|---|---|---|
| FALSE | FALSE | DPT | 9 |
| FALSE | FALSE | COL6A2 | 8 |
| FALSE | FALSE | LUM | 6 |
| FALSE | FALSE | THBS2 | 6 |
| FALSE | FALSE | ADAMTSL2 | 5 |
| FALSE | FALSE | AEBP1 | 5 |
| FALSE | FALSE | COL5A1 | 5 |
| FALSE | FALSE | LTBP2 | 5 |
| FALSE | TRUE | MMP2 | 5 |
| FALSE | FALSE | PHLDA1 | 5 |
| FALSE | FALSE | SMOC2 | 5 |

**Table 6:** LGLsrsi top 12 directly downstream genes

| In SenUMN? | In SenMayo? | **Other Genes** | **Total** |
|---|---|---|---|
| FALSE | FALSE | COL6A2 | 10 |
| FALSE | FALSE | COL4A2 | 8 |
| FALSE | FALSE | SPARC | 8 |
| FALSE | FALSE | THBS1 | 7 |
| FALSE | FALSE | TNXB | 7 |
| FALSE | FALSE | COL4A1 | 6 |
| FALSE | TRUE | MMP2 | 5 |
| FALSE | FALSE | ADAMTS2 | 4 |
| FALSE | FALSE | ATF3 | 4 |
| FALSE | FALSE | BGN | 4 |
| FALSE | FALSE | BHLHE40 | 4 |
| FALSE | FALSE | CFH | 4 |

There are some notable genes appearing on these lists. BOSSsrsi identified top gene, *IGFBP7,* is strongly implicated in chronic liver disease progression, operating as both an established tumor suppressor and a key inducer of senescence [42–44]. Additional upstream genes were heavily enriched for factors mediating extracellular matrix remodeling (*IGFPB7*, *VIM*, *MGP*, *DCN*), cellular stress responses (*JUNB*, *FOS*, *IGFBP3*, *TMSB4X*), and immune activation (*CD74*, *MIF*, *B2M*, *PTGDS*, *LYZ*). Downstream outputs were overwhelmingly dominated by structural ECM constituents and matrix-modulating enzymes, including multiple collagen isoforms (*COL6A1*/*2*/*3*), matrix metalloproteinase (*MMP2*), *TNXB*, *SPARC*, and *DCN*.

LGLsrsi identified a highly focused functional program with both upstream and downstream top genes overwhelmingly dominated by factors governing ECM structural assembly, organization, and turnover.

#### Pathway analysis

Due to the small number of genes identified by LGLsrsi, the ranked gene list pathway analysis was run only on the BOSSsrsi in order to increase power and reduce noise[45]. Gene Ontology (GO) biological process enrichment and Hallmark Gene Set Enrichment (GSE) analyses are shown in Figure 1. The results for the upstream variable sets are shown for GO and GSE in subfigures A and B, respectively. Downstream child pathway results are shown for GO and GSE in subfigures C and D, respectively.

**Figure 1.**
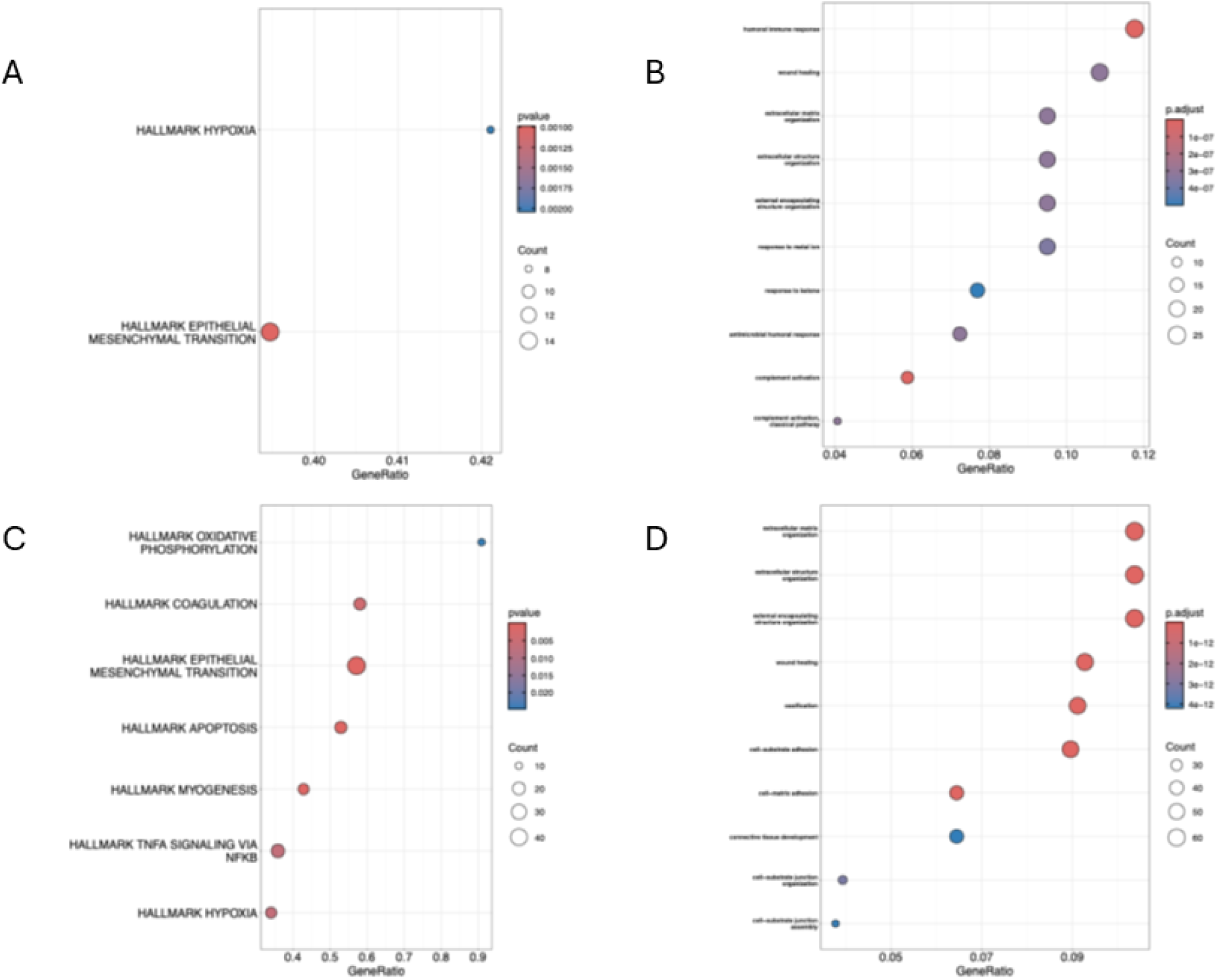
: ##

GO enrichment and Hallmark GSE analyses of parent genes revealed four primary overrepresented functional axes (Figures 1A and 1B). First, the niche exhibited robust activation of humoral immune responses and complement pathway cascades, reflecting adaptive immune response. Second, parent networks were heavily enriched for ECM programs, including *extracellular matrix organization*, *extracellular structure organization*, and *external encapsulating structure organization*. Third, we identified broad overrepresentation of cellular stress, specifically *response to oxidative stress*, *response to reactive oxygen species*, *response to hypoxia* (and *decreased oxygen levels*), *response to metal ion*, and *response to ketone*. Finally, we identified tissue repair cascades, including *wound healing* and *regeneration*.

The top three most significantly enriched downstream GO terms were strictly ECM-related, including *extracellular matrix organization*, *extracellular structure organization*, and *external encapsulating structure organization*, implicating fibrotic matrix deposition and structural stabilization represent a dominant phenotypic output of the senescent microenvironment (Figures 1C and 1D). Downstream genes also displayed enriched adaptive immune responses with signatures for *humoral immune response*, *complement activation*, and *humoral immune response mediated by circulating immunoglobulin* to reinforce the chronic immune engagement. Crucially, Hallmark GSE analysis showed that child genes were significantly enriched for *response to TNFa via NFKB response signaling pathway*, along with integrin and platelet-responsive pathways including *cell-substrate adhesion*, *cell-matrix adhesion*, and *integrin-mediated signaling pathways*.

## Discussion/conclusion

The work reported here is the first to perform spatial causal discovery analysis with well-described properties and guarantees of correctness on spatial omics data. The novel methods produced new insights into cellular senescence in human liver tissue. The analytical workflow involved numerous complex steps, including extensive cleaning, pre-processing, and data preparation, assessing the data’s viability for complex spatial causal modeling analysis, performing two different spatial causal discovery analyses on all data sets, extracting lists of genes that are modeled as upstream or downstream of senescence biomarker genes, and running pathway analyses on these genes.

### Based on the results from running these procedures, the original questions that guided this study can be answered as follows

A. *Are senescence biomarkers spatially autocorrelated?* Yes, there is significant spatial autocorrelation among the senescence biomarker genes used in this study. This is evident from the consistent significant rejection of the null hypothesis by Moran’s I tests.
B. *Do senescence biomarkers satisfy a Spatial Causal Markov Property?* Yes, the senescence biomarkers were found to be statistically independent, typically at a distance of 2 and occasionally at a distance of 3 Visium spots, when conditioning on the values of spatially separating spots. This helped ensure that a critical assumption of the causal discovery algorithms was satisfied.
C. *What molecular targets are upstream, and what molecular outcomes are downstream, of previously defined senescence biomarker genes?* The estimated causal models included a large number of genes that are not typically associated with senescence, since they are absent from the standard SenMayo and SenUMN gene lists.
D. *Can the novel gene-level spatial causal discovery approach introduced here produce valuable new insights into cellular senescence?* Yes, our novel algorithms and overall protocols used here provide new functional and causal perspectives on cellular senescence in human liver tissue. The combined integrin- and platelet-responsive pathways represent key mechanical and biochemical cascades known to directly trigger the activation of latent TGF-β within the remodeling extracellular matrix (Li et al. 2022). Both parent and child gene sets were enriched in the Hallmark epithelial-mesenchymal transition through GSE analysis. Together, the GO and Hallmark pathway profiles demonstrate that senescence-associated signaling integrates cellular stress inputs and immune responses to fuel a self-sustaining loop of integrin-mediated TGF-β activation and continuous matrix deposition. This suggests a research direction that should be investigated further.

### Limitations

Factors that limit the generalizability of the findings in this study include the following. The study was performed only on human liver tissue, so results may not apply to other tissue types or tissue from other animals. All results related to the ten identified senescence biomarkesr genes for human liver, however experts may prefer to use different sets of senescent biomarker genes. The number of tissue samples tested is only 11, so while each tissue sample contains a large number of spatial samples, biological variation that might occur between tissue samples may not be captured in its entirety.

### Future directions

Experimental validation of the modeled causal relationships reported here would be of great value. The approach used here should be extended to other animal models and other tissue types, and use additional gene sets to be tested as the target senescence biomarkers. Alternative discovery procedures could be applied and compared. A larger number of tissue samples could be analyzed, to increase robustness of findings to biological variation across tissue-samples.

## Biographical note

This paper reports the results of multiple complex analyses, including machine-learning-based spatial causal modeling, in the context of work conducted in two tissue mapping centers funded by the NIH senate consortium on cellular senescence.

## Acknowledgements

We thank the following groups, people, and entities for supporting this work: the SenNet Working group on Good Practices: Analytical Methods and Modeling (GP-AMM); the Clinical and Translational Science Institute (BLS Histology); the University of Minnesota Genomics Center; the resources and staff at the University of Minnesota University Imaging Centers (UIC) SCR_020997. The authors acknowledge the Minnesota Supercomputing Institute (MSI) at the University of Minnesota for providing resources that contributed to the research results reported within this paper.

## Appendix

This Appendix contains additional details.

## Appendix A: Spots remaining after BLADE, by tissue sample

278_D1 3,823
053_D1 4,387
289_D1 2,964
055_A1 4,431
289_A1 3,618
055_D1 4,316
278_A1 3,349
053_A1 4,572
110_A1 1,418
110_B1 1,335
110_C1 1,172

## Appendix B: Detailed results of testing Spatial Markov Property

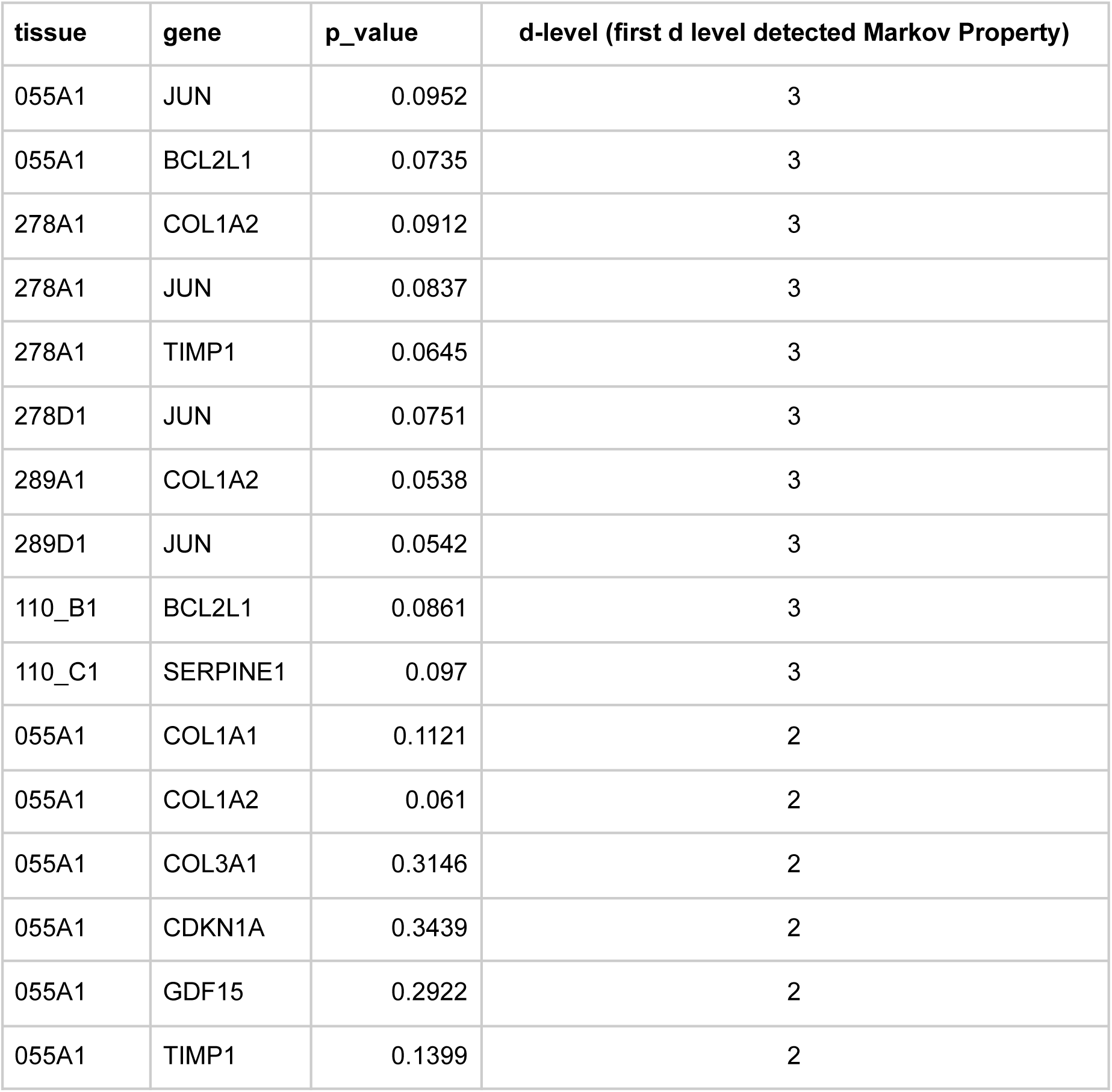
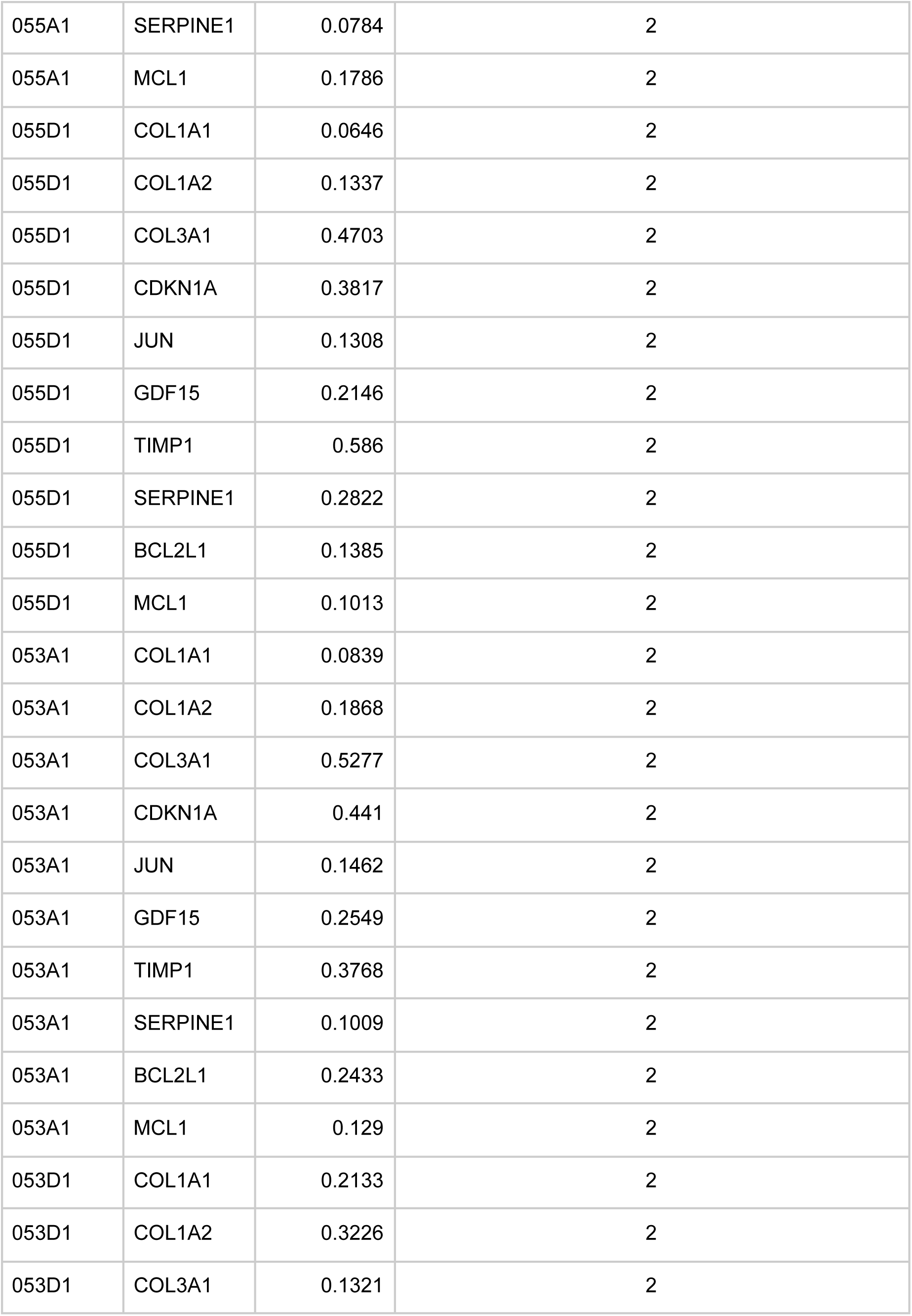
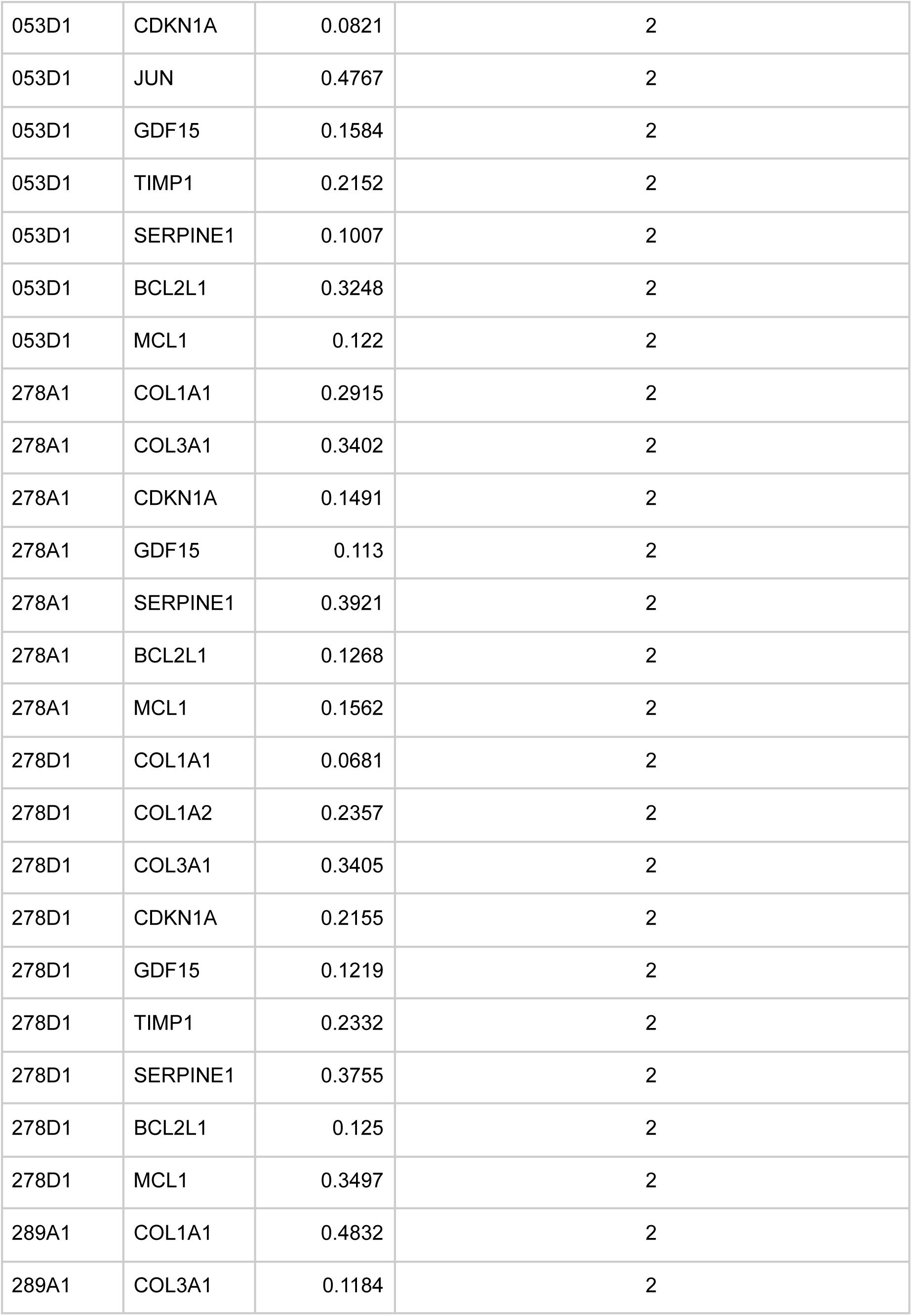
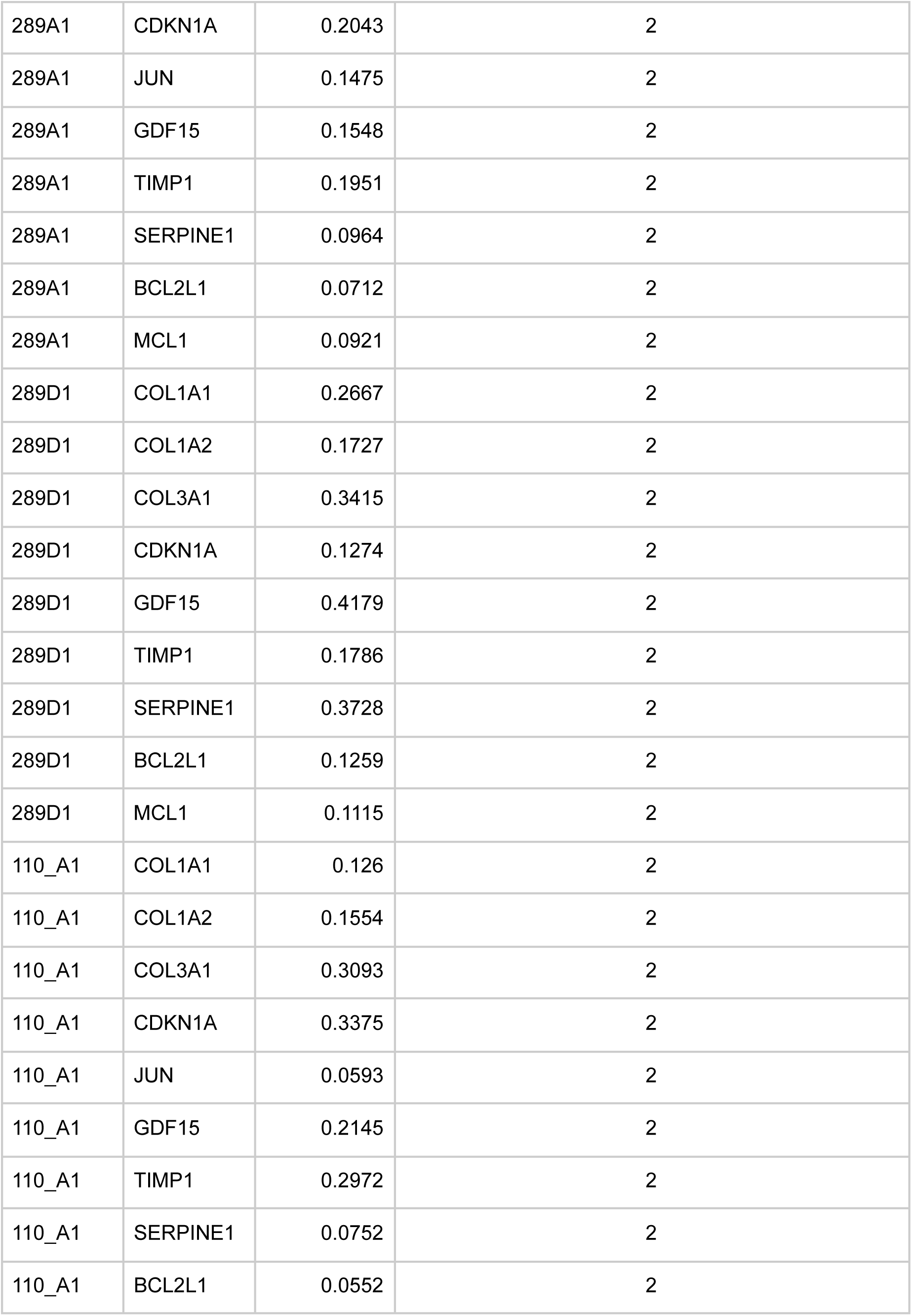
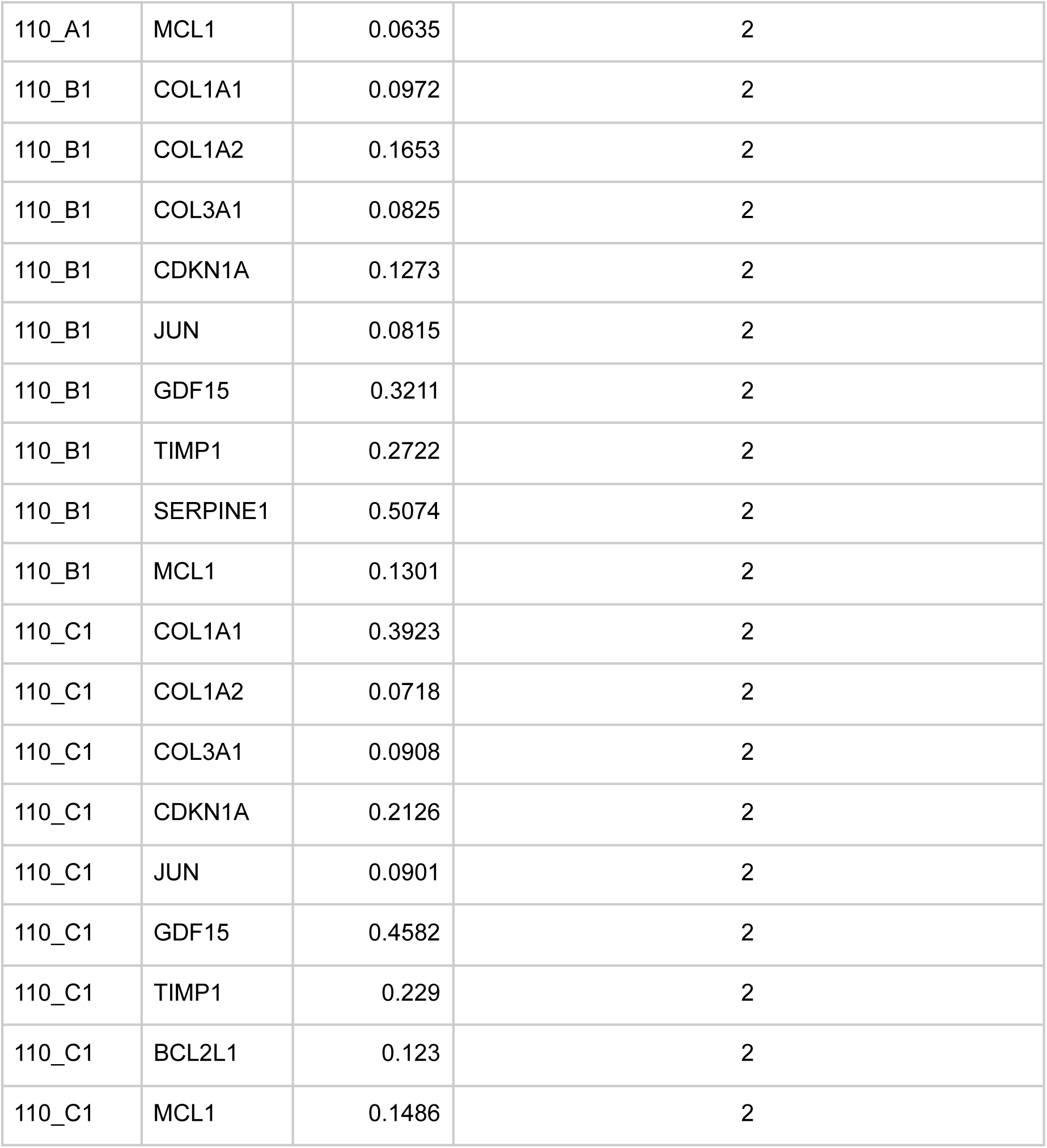

| **tissue** | **gene** | **p_value** | **d-level (first d level detected Markov Property)** |
|---|---|---|---|
| 055A1 | JUN | 0.0952 | 3 |
| 055A1 | BCL2L1 | 0.0735 | 3 |
| 278A1 | COL1A2 | 0.0912 | 3 |
| 278A1 | JUN | 0.0837 | 3 |
| 278A1 | TIMP1 | 0.0645 | 3 |
| 278D1 | JUN | 0.0751 | 3 |
| 289A1 | COL1A2 | 0.0538 | 3 |
| 289D1 | JUN | 0.0542 | 3 |
| 110_B1 | BCL2L1 | 0.0861 | 3 |
| 110_C1 | SERPINE1 | 0.097 | 3 |
| 055A1 | COL1A1 | 0.1121 | 2 |
| 055A1 | COL1A2 | 0.061 | 2 |
| 055A1 | COL3A1 | 0.3146 | 2 |
| 055A1 | CDKN1A | 0.3439 | 2 |
| 055A1 | GDF15 | 0.2922 | 2 |
| 055A1 | TIMP1 | 0.1399 | 2 |
| 055A1 | SERPINE1 | 0.0784 | 2 |
| 055A1 | MCL1 | 0.1786 | 2 |
| 055D1 | COL1A1 | 0.0646 | 2 |
| 055D1 | COL1A2 | 0.1337 | 2 |
| 055D1 | COL3A1 | 0.4703 | 2 |
| 055D1 | CDKN1A | 0.3817 | 2 |
| 055D1 | JUN | 0.1308 | 2 |
| 055D1 | GDF15 | 0.2146 | 2 |
| 055D1 | TIMP1 | 0.586 | 2 |
| 055D1 | SERPINE1 | 0.2822 | 2 |
| 055D1 | BCL2L1 | 0.1385 | 2 |
| 055D1 | MCL1 | 0.1013 | 2 |
| 053A1 | COL1A1 | 0.0839 | 2 |
| 053A1 | COL1A2 | 0.1868 | 2 |
| 053A1 | COL3A1 | 0.5277 | 2 |
| 053A1 | CDKN1A | 0.441 | 2 |
| 053A1 | JUN | 0.1462 | 2 |
| 053A1 | GDF15 | 0.2549 | 2 |
| 053A1 | TIMP1 | 0.3768 | 2 |
| 053A1 | SERPINE1 | 0.1009 | 2 |
| 053A1 | BCL2L1 | 0.2433 | 2 |
| 053A1 | MCL1 | 0.129 | 2 |
| 053D1 | COL1A1 | 0.2133 | 2 |
| 053D1 | COL1A2 | 0.3226 | 2 |
| 053D1 | COL3A1 | 0.1321 | 2 |
| 053D1 | CDKN1A | 0.0821 | 2 |
| 053D1 | JUN | 0.4767 | 2 |
| 053D1 | GDF15 | 0.1584 | 2 |
| 053D1 | TIMP1 | 0.2152 | 2 |
| 053D1 | SERPINE1 | 0.1007 | 2 |
| 053D1 | BCL2L1 | 0.3248 | 2 |
| 053D1 | MCL1 | 0.122 | 2 |
| 278A1 | COL1A1 | 0.2915 | 2 |
| 278A1 | COL3A1 | 0.3402 | 2 |
| 278A1 | CDKN1A | 0.1491 | 2 |
| 278A1 | GDF15 | 0.113 | 2 |
| 278A1 | SERPINE1 | 0.3921 | 2 |
| 278A1 | BCL2L1 | 0.1268 | 2 |
| 278A1 | MCL1 | 0.1562 | 2 |
| 278D1 | COL1A1 | 0.0681 | 2 |
| 278D1 | COL1A2 | 0.2357 | 2 |
| 278D1 | COL3A1 | 0.3405 | 2 |
| 278D1 | CDKN1A | 0.2155 | 2 |
| 278D1 | GDF15 | 0.1219 | 2 |
| 278D1 | TIMP1 | 0.2332 | 2 |
| 278D1 | SERPINE1 | 0.3755 | 2 |
| 278D1 | BCL2L1 | 0.125 | 2 |
| 278D1 | MCL1 | 0.3497 | 2 |
| 289A1 | COL1A1 | 0.4832 | 2 |
| 289A1 | COL3A1 | 0.1184 | 2 |
| 289A1 | CDKN1A | 0.2043 | 2 |
| 289A1 | JUN | 0.1475 | 2 |
| 289A1 | GDF15 | 0.1548 | 2 |
| 289A1 | TIMP1 | 0.1951 | 2 |
| 289A1 | SERPINE1 | 0.0964 | 2 |
| 289A1 | BCL2L1 | 0.0712 | 2 |
| 289A1 | MCL1 | 0.0921 | 2 |
| 289D1 | COL1A1 | 0.2667 | 2 |
| 289D1 | COL1A2 | 0.1727 | 2 |
| 289D1 | COL3A1 | 0.3415 | 2 |
| 289D1 | CDKN1A | 0.1274 | 2 |
| 289D1 | GDF15 | 0.4179 | 2 |
| 289D1 | TIMP1 | 0.1786 | 2 |
| 289D1 | SERPINE1 | 0.3728 | 2 |
| 289D1 | BCL2L1 | 0.1259 | 2 |
| 289D1 | MCL1 | 0.1115 | 2 |
| 110_A1 | COL1A1 | 0.126 | 2 |
| 110_A1 | COL1A2 | 0.1554 | 2 |
| 110_A1 | COL3A1 | 0.3093 | 2 |
| 110_A1 | CDKN1A | 0.3375 | 2 |
| 110_A1 | JUN | 0.0593 | 2 |
| 110_A1 | GDF15 | 0.2145 | 2 |
| 110_A1 | TIMP1 | 0.2972 | 2 |
| 110_A1 | SERPINE1 | 0.0752 | 2 |
| 110_A1 | BCL2L1 | 0.0552 | 2 |
| 110_A1 | MCL1 | 0.0635 | 2 |
| 110_B1 | COL1A1 | 0.0972 | 2 |
| 110_B1 | COL1A2 | 0.1653 | 2 |
| 110_B1 | COL3A1 | 0.0825 | 2 |
| 110_B1 | CDKN1A | 0.1273 | 2 |
| 110_B1 | JUN | 0.0815 | 2 |
| 110_B1 | GDF15 | 0.3211 | 2 |
| 110_B1 | TIMP1 | 0.2722 | 2 |
| 110_B1 | SERPINE1 | 0.5074 | 2 |
| 110_B1 | MCL1 | 0.1301 | 2 |
| 110_C1 | COL1A1 | 0.3923 | 2 |
| 110_C1 | COL1A2 | 0.0718 | 2 |
| 110_C1 | COL3A1 | 0.0908 | 2 |
| 110_C1 | CDKN1A | 0.2126 | 2 |
| 110_C1 | JUN | 0.0901 | 2 |
| 110_C1 | GDF15 | 0.4582 | 2 |
| 110_C1 | TIMP1 | 0.229 | 2 |
| 110_C1 | BCL2L1 | 0.123 | 2 |
| 110_C1 | MCL1 | 0.1486 | 2 |

